# The origin and evolution of acetylcholine signaling through AchRs in metazoans

**DOI:** 10.1101/424804

**Authors:** Dylan Z. Faltine-Gonzalez, Michael J Layden

## Abstract

**Background:** Acetylcholine is a cell signaling molecule that has been identified in plants, bacteria, and metazoans to play multiple roles in cells and as a neurotransmitter capable of exciting both neurons and muscle. While cell-cell communication activity has been reported in all phyla that have been investigated, its role as a neurotransmitter is less clear. Work within cnidarians implies that neurotransmitter activity emerged within or prior to the emergence of the cnidarian-bilaterian ancestor, but whether or not it is able to excite both muscles and neurons has not been addressed.

**Results:** To investigate the evolution of acetylcholine signaling we characterized the expression pattern of acetylcholine receptors (AchRs) and the neurotransmitter activity of acetylcholine in *Nematostella vectensis*. Expression patterns for 13 of the 21 known NvAchRs are consistent with acetylcholine acting as a cell signaling molecule and a neurotransmitter in neurons, muscles, or both. To dissect neurotransmitter activity we investigated the mechanism by which acetylcholine activates tentacular contractions in *Nematostella*. Tentacular contractions induced by application of acetylcholine are suppressed by inactivating voltage gated sodium channels with lidocaine indicating that acetylcholine specifically activates neurons in the tentacular contractile circuit.

**Conclusion:** Our results verify that acetylcholine’s neurotransmitter activity emerged prior to cnidarian-bilaterian divergence and that non-neuronal roles were likely retained in *Nematostella*. Additionally, we found no evidence to support a muscle activating role for acetylcholine indicating that its role in muscle excitability evolved during bilaterian evolution.

## INTRODUCTION

Cholinergic signaling is characterized by the release of the ligand acetylcholine from one cell, binding to and activating heteropenteramic acetylcholine receptors in responding cells (Dani, 2001). Acetylcholine and the enzymes necessary to synthesize it have been identified in bacteria, fungi, and all metazoan clades (Horiuchi et al., 2003; Kawashima et al., 2007; Joseph F. Ryan, 2014; Srivastava et al., 2008). However, acetylcholine receptors have not been identified outside of metazoans (Horiuchi et al., 2003). Within metazoans acetylcholine receptors have been described in bilaterians (vertebrates, echinoderms, insects, nematodes, and annelids, etc.) and cnidarians (sea anemones, corals and hydrozoans) (Angelini et al., 2004; Chapman et al., 2010; Lansdell & Millar, 2004; Noda et al., 1983.; Strader, Aglyamova, & Matz, 2018), cholinergic receptors have yet to be found in the basally branching placozoans, poriferans, and ctenophores (L. L. Moroz, 2015; Leonid L. Moroz et al., 2014; Leonid L. Moroz & Kohn, 2016; J. F. Ryan et al., 2013; Srivastava et al., 2008, 2010). There are two classes of acetylcholine receptors. The ionotropic nicotinic acetylcholine receptors (nAchRs) are ligand-gated ion channels that open in response to acetylcholine binding. The metabotropic muscarinic acetylcholine receptors (mAchRs) are G-coupled proteins that activate the associated G-protein signaling cascade that elicits a cellular response when acetylcholine is bound (Dani & Bertrand, 2007; Eglen, 2006). To date muscarinic receptors are exclusively found in bilaterian animals, whereas previous reports identified nicotinic receptors in multiple cnidarian species (Anctil, 2009; Chapman et al., 2010; Kass-Simon & Pierobon, 2007). These observations argue that cholinergic signaling through nicotinic acetylcholine receptors evolved prior to cnidarian-bilaterian divergence, but do not provide insights about the ancestral roles for cholinergic signaling.

All work to characterize the function of cholinergic receptors has been carried out in bilaterian species thus far. Both nicotinic and muscarinic receptors are best known for their role as neurotransmitters involved in neuronal communication at chemical synapses (Brown, 2010; Dani & Bertrand, 2007). In neurons acetylcholine activates or inhibits action potentials in receiving cells (Dani, 2001). Both nicotinic and muscarinic receptors also function to activate muscle contractions and as regulators of cell-cell signaling that results in changes in gene expression, proliferation, and apoptosis in responding cells (Wessler & Kirkpatrick, 2008). The most notable difference between muscarinic and nicotinic receptors is that nAchRs activate muscle contractions primarily in skeletal muscle whereas mAchRs primarily regulate smooth muscle contractions (Dani & Bertrand, 2007, Eglen, 2006). Non-neuronal acetylcholine responses have been described within all metazoans as well as fungi, plants, and bacteria (Horiuchi et al., 2003; Kawashima et al., 2007). Within plants acetylcholine has been found to regulate growth, cellular differentiation, water homeostasis, and photosynthesis (Kawashima et al., 2007). While non-neuronal acetylcholine within mammalian cells has been implicated in signal transduction, cell-cell contact, proliferation, differentiation, and cell migration (Wessler & Kirkpatrick, 2008). As acetylcholine has ancient roles regulating multiple aspects of biology in single and multicellular organisms, cholinergic signaling likely emerged as a mechanism to improve the resolution of acetylcholine activity to regulate the biology of multicellular animals through cell-cell communication, and that it was later co-opted to regulate neuronal signaling and muscle contractions. However, because data investigating acetylcholine receptors outside of the bilaterians is sparse, it is not clear when the neural and muscular functions of acetylcholine signaling evolved. Because Cnidaria is the sister taxon to Bilateria and they are the only other metazoan clade where AchRs have been identified, investigating the role of acetylcholine in these animals is critical to understand the origin and evolution of acetylcholine signaling (Anctil, 2009; Chapman et al., 2010; Dunn et al., 2008; Srivastava et al., 2008).

Characterization of acetylcholine biology within cnidarians has focused on genomic studies to identify individual components of cholinergic signaling and a number of pharmacological studies aimed at determining its possible role in neuronal signaling. (Anderson & Trapido-Rosenthal, 2009; Denker, Chatonnet, & Rabet, 2008; rev. Kass-Simon & Pierobon, 2007; Oren, Brikner, Appelbaum, & Levy, 2014; Pierobon, 2012; Sebé-Pedrós et al., 2018; Takahashi & Hamaue, 2010). The earliest studies isolated acetylcholine from cnidarian tissue and showed that application of acetylcholine induced tentacular contractions in both anthozoans and hydrozoan species suggesting acetylcholine is a conserved regulator of tentacular contraction in cnidarians (Mendes & De Freitas, 1984; Singer, 1964). The availability of genomic data has allowed genes necessary for cholinergic signaling to be identified in a greater number of cnidarians species (Anctil, 2009; Oren et al., 2014; Pierobon, 2012). Spatiotemporal expression analysis using mRNA *in situ* hybridization identified that acetylcholinesterase, an enzyme necessary to rapidly metabolize acetylcholine, was broadly expressed in endodermal tissue of both *Hydra* and the sea anemone *Nematostella*, resembling a more non-neuronal phenotype for acetylcholine (Pierobon 2007, Oren et al.2014). The culmination of this data while vast has failed to describe expression of the receptors necessary to bind acetylcholine *in vivo*, which is essential to understand the putative ancestral roles of acetylcholine signaling. Recent work in *Nematostella* carried out a series of single cell RNA sequencing experiments that identified expression of nAchRs in neuronal, muscle cells, and gland/secretory cells as well as RNA sequencing of the apical tuft which has identified nAchRs expression (Sebé-Pedrós et al., 2018; Sinigaglia, Busengdal, Lerner, Oliveri, & Rentzsch, 2015). Taken together the data from cnidarian species suggests that of acetylcholine receptor function in neurons, muscle contractions, and cell-cell signaling all arose near the emergence of the AchRs in animal evolution, but functional studies are needed to confirm this hypothesis.

Here we characterized the roles of the acetylcholine receptors in the anthozoan cnidarian sea anemone *Nematostella vectensis*. Previous work has identified three choline acetyltransferases, five copies of acetylcholinesterase, and 12 nicotinic acetylcholine receptors encoded within the annotated genome (Anctil, 2009). Research into the role of acetylcholine within *Nematostella* identified the localization of one of the three acetylcholinesterase genes identified and determined it’s role to be non-neuronal due to its expression within non-neuronal endodermal tissue (Oren et al., 2014). Single cell RNA sequencing within juvenile polyps identified nicotinic acetylcholine receptor expression within both neural cells as well as tentacular and longitudinal muscle cells (Sebé-Pedrós et al., 2018). We identified nine additional nAchRs in the *Nematostella* genome, describe the spatiotemporal expression patterns for 13 of the 21 nAchRs, and carried out pharmacological studies to better understand the role acetylcholine plays in regulating tentacular contractions. Expression of AchRs ranged from ubiquitous to expressed in few scattered cells that resemble the salt and pepper expression exhibited by known neuronal markers (Layden, Boekhout, & Martindale, 2012; Rentzsch, Layden, & Manuel, 2017). Tentacular expression was consistent with being expressed in neurons, muscles, or both. We also found expression within the pharynx and the apical tuft, which are non-neuronal expression patterns. Pharmacological experiments suggest that tentacular contractions are likely regulated by acetylcholine activity in neurons rather than muscle cells. The culmination of the data presented here suggests that utilization that acetylcholine receptors functioned in both neuronal and non-neuronal roles in the cnidarian-bilaterian ancestor, and suggest that acetylcholine’s direct role in muscle contractions likely evolved within the bilaterian lineage.

## METHODS

### Phylogenetic Analysis

Bilaterian protein sequences were found using keywords (nicotinic acetylcholine receptor, muscarinic acetylcholine receptor, GABAnergic receptor, and glycine receptor) to identify sequences within multiple genomic databases <u>mouse genome informatics, flybase, wormbase</u> and <u>zfin</u>. *Nematostella* coding sequences were then collected from the US Department of Energy Joint Genome Institute, *Nematostella* genome site (https://genome.jgi.doe.gov/pages/search-for-genes.jsf?organism=Nemve1) by blasting bilaterian nicotinic acetylcholine receptor sequences against the *Nematostella* genome. The top ten *Nematostella* BLAST hits were then selected and uploaded to ApE v2.0.47 where they were translated. The largest open reading frame identified in the *Nematostella* coding sequences were then used for protein alignment (Paradis, Claude, & Strimmer, 2004). All protein sequences were uploaded into MEGA v7.0.25 software and a MUSCLE alignment was performed (Kumar, Stecher, & Tamura, 2016). *Nematostella* sequences missing the key residues for nicotine binding were deleted before phylogenetic analysis was performed. Muscarinic acetylcholine receptors were used to determine specificity of clustering to nicotinic acetylcholine receptors. GABA and glycine receptors were used as the outgroup for the nicotinic acetylcholine receptors due to their shared ancestral origins (Anctil, 2009). Phylogenetic evolution models were determined using the protein model function on MEGA v7.0.25 software (Kumar et al., 2016).

#### Animal care

Adult *Nematostella* were maintained at Lehigh University in 17°C incubators. Culture and spawning of animals were performed according to previously published protocols (Fritzenwanker & Technau, 2002). Fertilized embryos were raised at 22°C until fixation at the required stage.

### *In situ* hybridization and Immunohistochemistry

Embryo fixation, RNA probe synthesis, in situ hybridization, and immunohistochemistry protocols were performed as previously described (Havrilak et al., 2017; Wolenski, Layden, Martindale, Gilmore, & Finnerty, 2013). Juvenile polyps were fixed for two hours as previously described (Steinmetz, Aman, Kraus, & Technau, 2017).

### Drug Treatments

Two week old, one, two, and three month old juvenile polyps were placed into a 5mm petri dish and time series was performed for five minutes. At the two minute mark 1mL of 50mM acetylcholine-chloride (Sigma-Aldrich Cat.#A6625) or nicotine was added to 4mLs of 1/3X artificial sea water to make a final concentration of 10mM. To block potential acetylcholine receptors, juvenile polyps were treated with 50μL of 50mM mecamylamine hydrochloride (Sigma-Aldrich Cat.#M9020) added to 5mLs of 1/3X artificial seawater for a final concentration of 500μM ten minutes prior to treatment with acetylcholine-chloride. 500μL of 100mM Lidocaine (Sigma-Aldrich Cat.#L7757) was added to 5mLs of 1/3X artificial sea water for a final concentration of 10mM was used to block sodium channels before treatment with 10mM acetylcholine. Controls were performed by replacing pharmacological drugs with 1/3X artificial seawater.

### Imaging

Images for *in situ* hybridization were taken on a Nikon NTi with a Nikon DS-Ri2 color camera using the Nikon elements software. Time series were performed on a Nikon Eclipse E1000 with a Nikon and analyzed on the Nikon elements software. Time series images were analyzed on the Fiji software v.2.0.0-rc-54/1.51g (Schindelin et al., 2012). All images were edited using Illustrator and Photoshop (Adobe Inc.).

## RESULTS

### Identification of acetylcholine receptors in Nematostella vectensis

Our BLAST searches of updated genomic sequence data suggested that ten previously unknown putative acetylcholine receptors are encoded in the *Nematostella* genome. We excluded one of the ten putative acetylcholine receptors, because it lacked the conserved Y190, Y198, C192, and C193 motif necessary to form the acetylcholine binding pocket (Supplementary figure 1)(Anctil, 2009; Unwin, 2005). To determine if the remaining nine putative receptors were cholinergic we performed a phylogenetic analysis. All 21 receptors grouped with the acetylcholine receptors with 100% bootstrap support using Maximum Likelihood with the LG model and GABA/Glycine receptors as the outgroup (Figure 1, blue box). Consistent with previous findings, all *Nematostella* receptors clustered within the nicotinic class of acetylcholine receptors, which is further supported by the fact that none of the *Nematostella* sequences encode the Q207(M3)/Q163(M2) and L204 motif that is necessary for G-coupled signaling characteristic of muscarinic receptors (Figure 1, Supplemental Figure 2)(Kruse et al., 2012). We conclude that we identified seven previously undescribed acetylcholine receptors and that *Nematostella* has 21 total nicotinic acetylcholine receptors.

**Figure 1:**
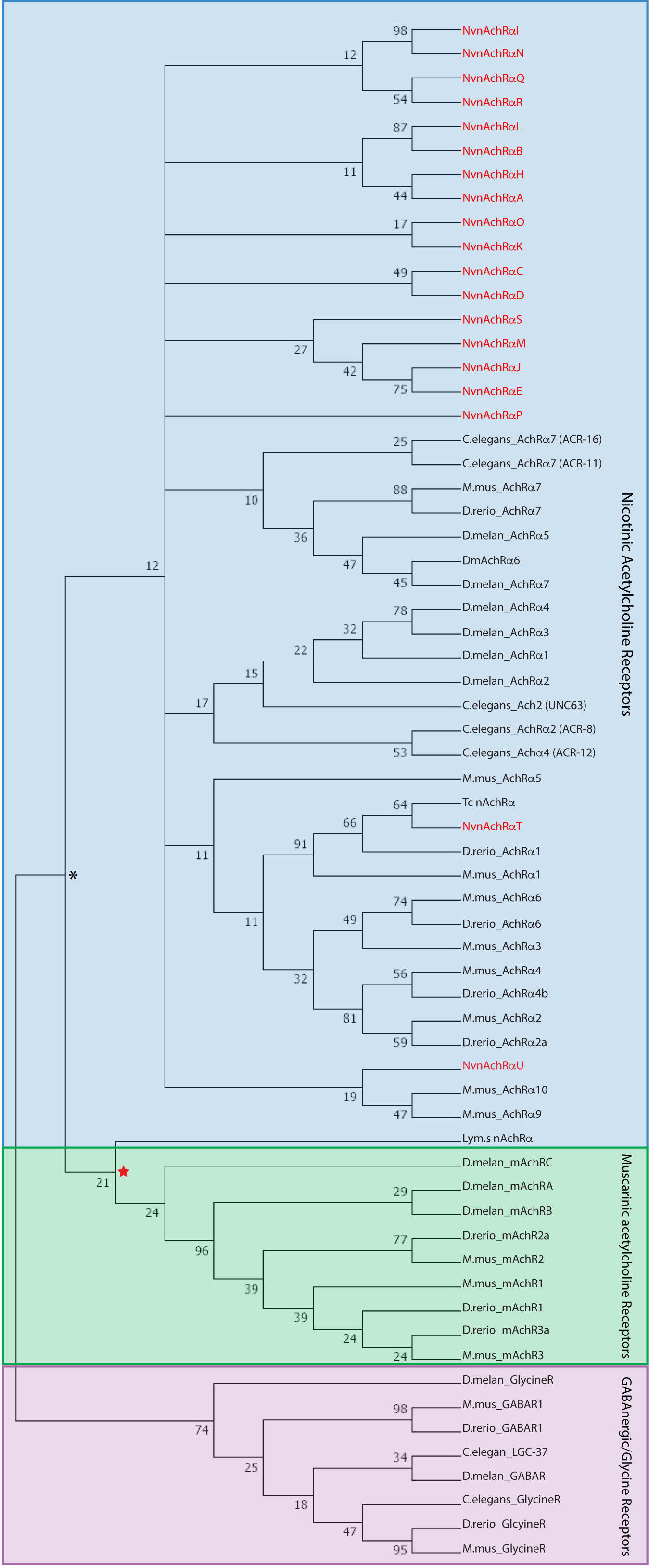
Phylogenetic analysis of *Nematostella* AchRs. Maximum likelihood tree determining the relationship of potential *Nematostella* nicotinic acetylcholine receptors (text in red) with bilaterian nicotinic acetylcholine receptors (Blue Box) and muscarinic acetylcholine receptors (Green Box). The red star indicates the node where nicotinic and muscarinic receptors diverged. Protein sequences were trimmed to consist of the acetylcholine binding pocket and the resides for nicotinic binding. Glycine and GABA receptors were used as the outgroup (Red Box) For alignment see supplemental table 1. Numbers indicate the bootstrap values calculated at each node and the asterisk indicates 100% bootstrap values.

Our phylogenetic analysis did not suggest that any single *Nematostella* cholinergic receptors was a definitive ortholog of previously described bilaterian subunits. Three publications previously described *Nematostella* cholinergic genes, but there is little consensus on the naming of these genes (Oren et al., 2014; Sebé-Pedrós et al., 2018; Sinigaglia, Busengdal, Lerner, Oliveri, & Rentzsch, 2015b). We have named the 21 genes *NvnAchRα A- NvnAchRαU* (Supplemental Table 1). So as not to confuse them with the characterized bilaterian subunits that are classified using a number to define orthologous receptors.

### Determining spatiotemporal expression of acetylcholine receptors

To determine putative roles for the acetylcholine in *Nematostella* we characterized the spatiotemporal expression pattern of the receptors by mRNA *in situ* hybridization. We were able to clone and generate *in situ* probes for 13 of the 21 receptors all of which were expressed at some point during development (Figure 2). The earliest expressed were *NvnAchRαI* and *NvnAchRαF*. *NvnAchRαF* is expressed in salt and pepper pattern within gastrula and *NvnAchRαF* was expressed ubiquitously in gastrula and then lost in planula stages (Figure 2AK and AL). The ubiquitous expression of *NvnAchRαF* is consistent with a role in cell-cell signaling, consistent with previously described roles for cholinergic signaling in epithelia (Wessler & Kirkpatrick, 2008). While the gastrula salt and pepper expression of *NvnAchRαI* is consistent of a neuronal phenotype and a neuronal role found in bilaterians (Dani, 2001; Layden et al., 2012; Nakanishi, Renfer, Technau, & Rentzsch, 2012; Richards & Rentzsch, 2015a). At larval stages we detected a number of distinct expression patterns. Three (*NvnAchRαI*, *NvnAchRαH*, and *NvnAchRαG*) were expressed in a small subset of scattered cells in the pharyngeal endoderm (Figure 2A-H). Pharyngeal expression was first detected in planula stages (Figure 2B-C, F-G, J-K), and it was maintained and into the juvenile polyp stage (Figure 2D, H, L). The scattered cell pattern is characteristic of but not exclusively consistent with expression of known neuronal markers in *Nematostella* (Layden et al., 2012; Nakanishi et al., 2012; Richards & Rentzsch, 2014). Two genes, *NvnAchRαJ* and *NvnAchRαK*, are expressed in the apical tuft ectoderm of planula larvae, confirming the published expression patterns of *NvnAchRαJ* (199721/ao145) and *NvnAchRαK* (110265/ao19) (Figure 2M-T) (Sinigaglia et al., 2015). This apical tuft expression was only maintained during the planula stages and is subsequently lost in juvenile polyps (Figure 2P, T). Acetylcholine signaling has been linked to metamorphosis in bivalve free swimming larvae and the apical tuft organ is linked to settlement and metamorphosis in some corals, suggesting that the apical tuft expression may indicate a conserved role for acetylcholine in anthozoan cnidarian metamorphosis (Sánchez-Lazo, Martínez-Pita, Young, Alfaro, 2012; Strader et al., 2018).

**Figure 2:**
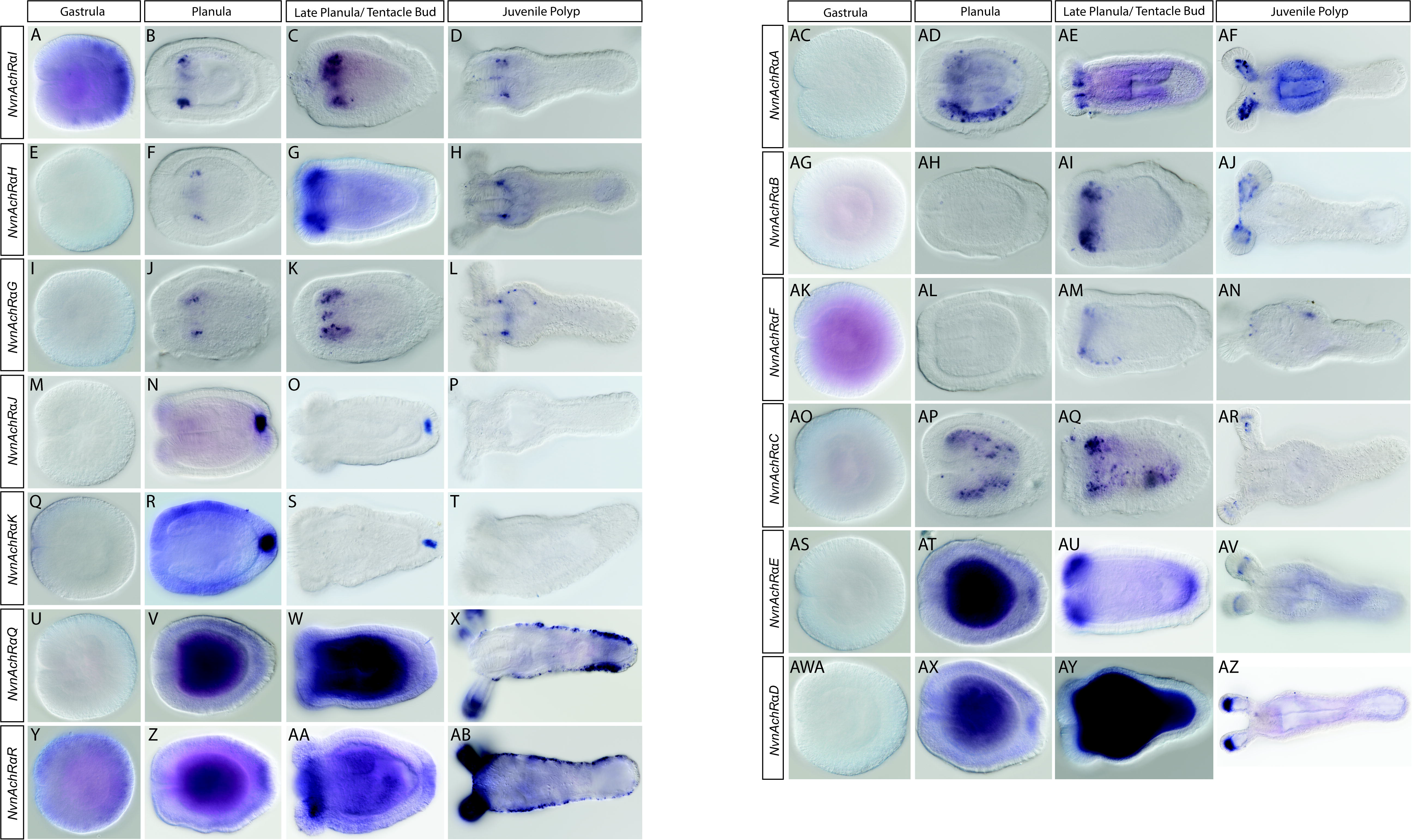
Expression patterns of *Nematostella* nicotinic acetylcholine receptor genes. *NvnAchRαA* expression is salt and pepper within the endoderm and then restricted towards the tentacle tips at the tentacle bud stage (B, C). This expression then becomes isolated towards the tentacular endoderm (D). *NvnAchRαB* expression starts at the tentacle bud stage found in the endoderm within the developing tentacles (F, G). This continues in the juvenile polyp with expression observed within endoderm of the tentacles (H). *NvnAchRαF* is expressed at the tentacle bud stage in the endodermal tissue where the tentacles will develop with no expression before that (K). By the juvenile polyp stage expression in a few cells can be found at the distal ends of the tentacles (L). *NvnAchRαC* has a salt and pepper expression pattern within the planula endoderm (N). This expression becomes restricted to the tentacle tips at the tentacle bud stage and then becomes restricted to the proximal endoderm and ectoderm of the tentacles (O,P). *NvnAchRαE* expression starts at the planula stage and continues into the tentacle bud stage with expression at the site where the tentacle tips will develop (R,S). Expression then becomes isolated to the ectodermal region just below the most proximal end of the tentacles (T). Lastly *NvnAchRαD* is expressed within the ectoderm surrounding the tissue developing into the tentacles. This expression is then isolated to the most proximal end of the tentacles (X). *NvnAchRαI*. *NvnAchRαH*, and *NvnAchRαG* are expressed within the pharyngeal ectoderm starting from the planula stage to the juvenile polyp stage (AG-AR). *NvnAchRαM* and *NvnAchRαO* are both expressed in the apical tuft of the planula and tentacle bud and lost in the juvenile polyps (Y-AF). All images have the oral end directed to the left.

Six nicotinic acetylcholine receptors displayed tentacular expression patterns (Figure 2AC-AZ). *NvnAchRαA* and *NvnAchRαC* are first expressed in a salt and pepper expression pattern, typical of neuronal genes, within the planula endoderm (Figure 2 AD, AP). At the tentacle bud stage expression of both becomes restricted to the endodermal tissue beneath the tentacle tips (Figure 2AE, AQ). At the juvenile polyp stage the salt and pepper expression of *NvnAchRα*A is observed along the length of the tentacle endoderm (Figure 2AF), while *NvnAchRα*C is restricted to the distal ends of the tentacle (Figure 2AR). The other four acetylcholine receptors are expressed within the endoderm below the developing tentacles (Figure2 AI, AM, AU, AQ). At the juvenile polyp stage *NvnAchRαB* is expressed in a salt and pepper pattern within the tentacular endoderm (Figure 2AJ). *NvnAchRαE* expression is restricted to the distal end of the tentacular endoderm (Figure 2AV). *NvnAchRαD* is expressed within the distal ends of the tentacular endoderm (Figure 2 AZ). Lastly, *NvnAchR αF* is expressed within the very proximal base of the tentacles (Figure 2AN).

The expression patterns suggest multiple roles of acetylcholine signaling within *Nematostella*. The ubiquitous expression patterns found within *NvnAchRαF* are reminiscent of a cell to cell signaling that could be occurring within *Nematostella*. The salt and pepper expression found within the tentacles suggests expression within a neuronal subtype localized to the tentacle tips. The tentacular expression within the tentacular endoderm along the length of the tentacle is likely a mixture of both could be tentacular neurons and myoepithelial cells both of which were recently shown to express a similar suite of *NvAchR’s* by single cell RNA sequencing data (Sebé-Pedrós et al., 2018). Bilaterian AchRs function within multiple non-neuronal cell types, neuronal cells, and muscles cells. Our expression data argue that all three roles described in bilaterians are likely present within the cnidarians.

### Acetylcholine treatment induces tentacle specific contractions

Previous reports in one hydrozoan and one anthozoan species suggested that acetylcholine is capable of inducing tentacular contractions (Kass-Simon & Pierobon, 2007). To determine if acetylcholine mediated tentacular contraction is conserved in *Nematostella* we treated 3 month old adult polyps with 10mM acetylcholine. Treatment with acetylcholine resulted in rapid contraction of tentacles in 96%, 100%, 85% (Figure 3A-C; Figure 4D-F; Supplemental video 1 and 2) of animals compared to the 14% and 4% of observed tentacles that contracted in water treated controls (Figure 3 A-C, P; Figure 4 A-C,P, Supplemental Video 6). The tentacles relaxed after acetylcholine treatment, presumably because of acetylcholinesterase activity (Figure 3C; Supplemental video 1). To confirm specificity of this contractile response to acetylcholine we treated juvenile polyps with a broad non-competitive antagonist of acetylcholine receptors, mecamylamine. Animals were treated with 500μM mecamylamine prior to treatment with acetylcholine. This pretreatment was sufficient to block acetylcholine induced tentacle contractions (Supplemental Video 3). Only 18% of mecamylamine treated animals contracted in response to acetylcholine, which resembled controls treated with water (Figure3 G-I, P, Supplemental Video 3). To determine if these induced tentacle contractions were due to nicotinic acetylcholine receptors we treated juvenile polyps with 10mM nicotine a known agonist for nicotinic acetylcholine receptors. Nicotine induced a more robust (100%) tentacle contraction than what was observed during the acetylcholine treatments (Figure 3 D-F, P; Figure 4J-L, P, Supplemental Video 4). Treatment with nicotine resulted in complete loss of the subsequent relaxation presumed to be due to the inability of acetylcholinesterase to metabolize nicotine and/or the fact that nicotine binds cholinergic receptors with higher affinity than acetylcholine (Jadey, Purohit, & Auerbach, 2013)(Figure 3F). To determine if tentacular contractions are induced by acetylcholine at all polyp stages we repeated our experiments in two week old juvenile polyps, and one and two month old polyps (Figure 3Q). The percentage of animals with tentacular contractions were ≥ 78% at all stages indicating that acetylcholine regulates tentacular contractions at all life stages. This is consistent with our previous observations that juvenile polyp nervous systems resemble adult nervous systems, but have fewer total neurons (Havrilak et al., 2017). We conclude that acetylcholine induces tentacular contractions in *Nematostella*, and that acetylcholine mediated tentacular contraction is likely a conserved trait in cnidarian polyps.

**Figure 3:**
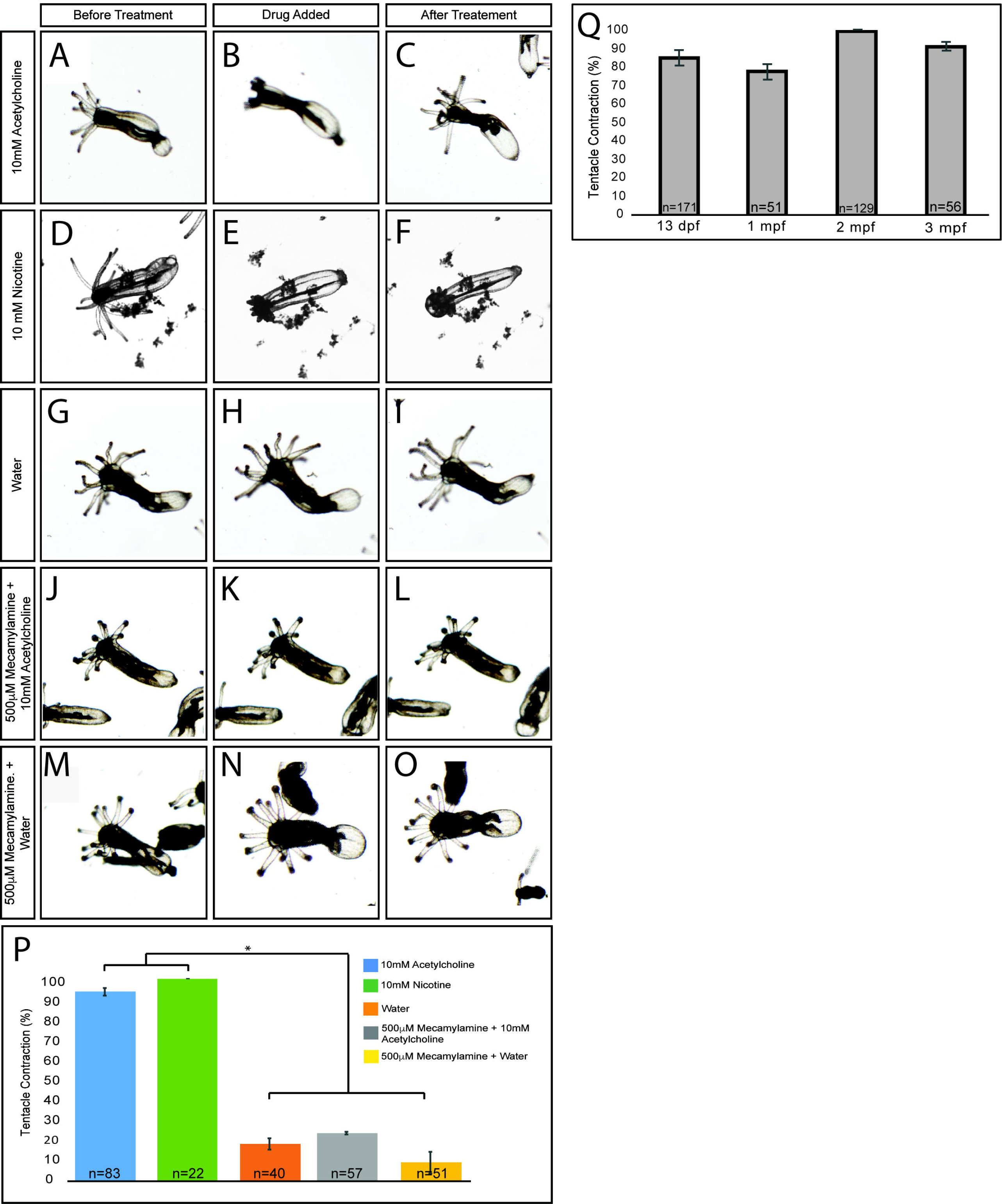
Pharmacological analysis of acetylcholine’s role in tentacular contractions. Treatment with 10mM acetylcholine resulted in tentacle contraction in 93% of the juvenile polyps tested (A and B, P). These tentacles then relaxed after the initial acetylcholine induced contraction (C). Treatment with 10mM nicotine resulted in tentacle contraction in 100% of the juvenile polyps tested with no relaxation observed after treatment (D-F and P). Pretreatment with 500 μM mecamylamine prior to treatment with 10mM acetylcholine resulted in loss of induced tentacle contractions in 22% of juvenile polyps (J-L and P). This blocked response to acetylcholine is similar that observed when treated with water (p<0.05) (J-O,P). Contractile response to endogenous acetylcholine is maintained from two weeks post fertilization to three months post fertilization (p>0.05)(Q). Quantifications comparing the contraction of tentacles is summarized in P with asterisks indicating significant difference, p<0.05. All images have the oral end directed to the left of the figure.

**Figure 4:**
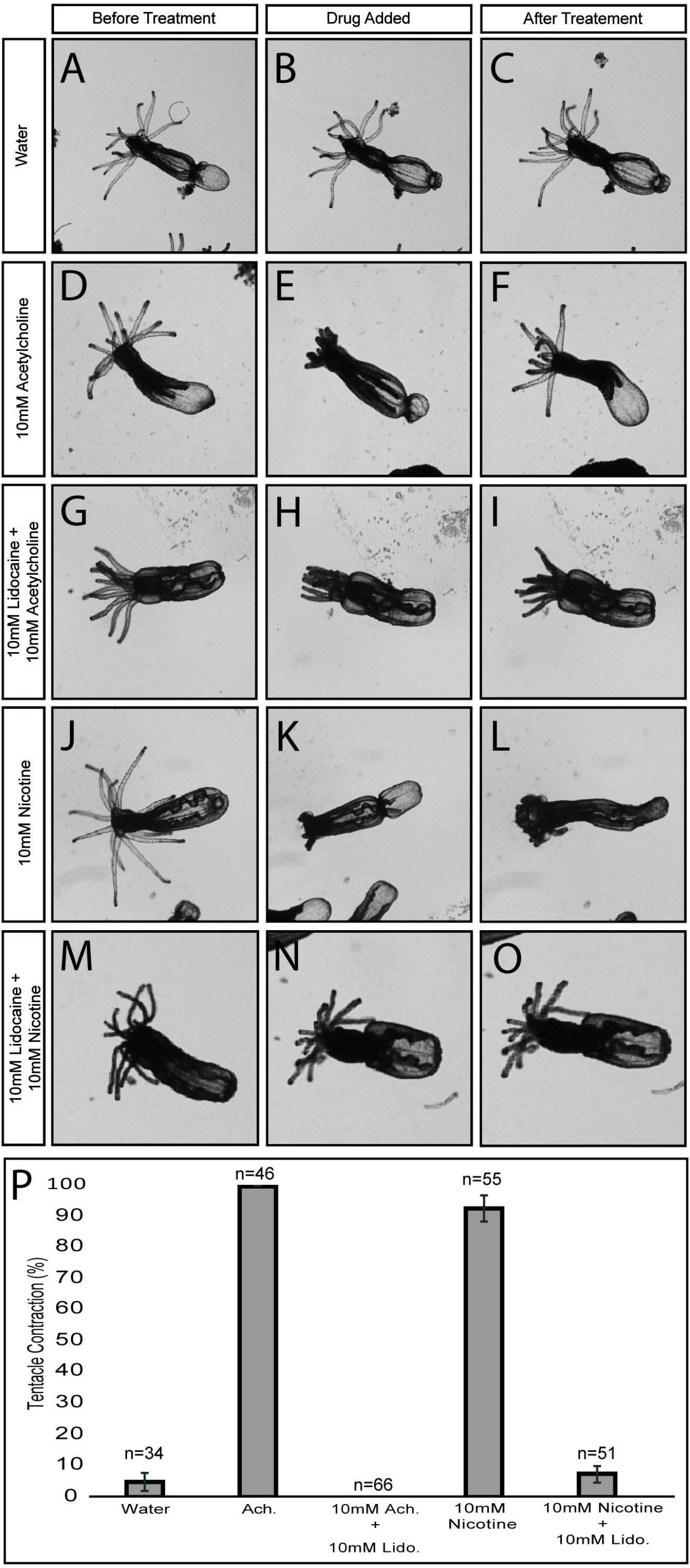
Lidocaine suppresses acetylcholine mediated tentacular contractions. Pretreatment with 10mM lidocaine resulted in loss of acetylcholine induced tentacle contraction in all the juvenile polyps tested (H and P). This was repeated in polyps treated with 10mM lidocaine and then nicotine (N and P). These losses are significantly different then when juvenile polyps are treated with acetylcholine (E) or nicotine (k) alone, p<0.05%. Quantifications comparing the contraction of tentacles are summarized in P with asterisks indicating significant difference, p<0.05. All images have the oral end directed to the left of the figure.

### Lidocaine treatment implies acetylcholine induced muscle contractions are due to neuronal activation

Both muscle cells and neurons are reported to express acetylcholine receptors (Figure 2A and B) (Sebé-Pedrós et al., 2018). To determine if acetylcholine induced muscle contractions are more likely due to acetylcholine directly inducing myoepithelial cell contraction or due to activation of a cholinergic neuron upstream of tentacular muscles we treated animals with lidocaine prior to application of acetylcholine. Lidocaine is a broad inhibitor of voltage gated sodium channels. Inhibition of voltage gated sodium channels suppresses neuronal action potentials (Sheets & Hanck, 2003). If lidocaine treatment inhibited acetylcholine’s ability to induce tentacle contractions, it would suggest that acetylcholine is activating neurons upstream of the myoepithelial cells rather than directly activating the muscle cell. Pretreatment with 10mM lidocaine resulted in a loss of acetylcholine induced tentacle contractions. No animals contracted in the presence lidocaine compared to 100% contracted animals in the acetylcholine treated group (Figure 4G-I, P)(Supplementary videos 5). This loss of tentacular contraction was repeated in nicotine treated animals as well (Figure 4M-O, P). The ability of lidocaine to block both acetylcholine and nicotine induced tentacle contractions suggests that application of acetylcholine activates a neuron(s) upstream of tentacular muscles rather than directly inducing muscular contraction.

## Discussion

### Neuronal activity of acetylcholine modulates muscular contractions in Nematostella

Acetylcholine has defined roles at neuronal synapses and neuromuscular junctions in bilaterian animals. In multiple cnidarian species treatment with acetylcholine induces tentacular contractions (Kass-Simon & Pierobon, 2007; Singer, 1964). Our data and single cell RNA sequencing data identified *NvnAchRαD*, *NvnAchRαE*, *NvnAchRαR* expression in tentacles of juvenile polyps in patterns consistent with being expressed in both neurons and muscle (Figure 2AB, AF, AJ). (Sebé-Pedrós et al., 2018). Thus, expression patterns alone are not capable of distinguishing whether or not acetylcholine activates neurons, muscles, or both cell types in *Nematostella*. To determine if acetylcholine could activate muscles directly, we first confirmed that muscle contractions are likely due to the specific activity of nicotinic acetylcholine receptors using a series of pharmacological treatments with the agonists acetylcholine and nicotine, and the receptor antagonist mecamylamine (Figures 3,4; Supplemental video 1 and 2). We then reasoned that if acetylcholine activates muscle cells that inhibition of neuronal activity using the voltage gated sodium channel blocker lidocaine should have no impact on tentacular contractions following acetylcholine treatment. Conversely, if lidocaine treatment inhibits tentacular contractions, then acetylcholine is likely activating a neuron upstream in the tentacular muscle. Lidocaine blocked tentacular contractions for both nicotine and acetylcholine treated animals (Figure 4). This suggests that neurons, not myoepithelial cells, are activated to induce tentacular contractions when acetylcholine is applied. Our combined pharmacological and gene expression data suggest that the cnidarian-bilaterian ancestor likely used acetylcholine as a neuronal modulator. It would also imply that the neuronal role of acetylcholine likely emerged shortly after or coincidently with the emergence of acetylcholine receptors, because no receptors have been identified in more basally branching metazoans (Chapman et al., 2010; Joseph F. Ryan et al., 2013; Srivastava et al., 2008, 2010). Our data also implies that acetylcholine activation of muscle cells likely evolved in bilaterians after the cnidarian-bilaterian split. This conclusion is supported by the observation that acetylcholine functioning to induce muscle contractions is not widespread within bilaterian animals (Ahmed & Ali, 2016; Goldman & Staple, 1989).

### Non-neuronal roles of acetylcholine within Nematostella

We identified the localization of a 13 out of the 21 nicotinic acetylcholine receptors found within the *Nematostella* genome. Only a small subset showed a neural or muscle like expression pattern (Figure 2). Genes that appear to be non-neuronal and non-muscle tend to be broadly expressed in one or more tissues (Figure 2). The broad expression patterns are consistent those cells being able to respond to acetylcholine signaling, but predicting the cellular response is difficult. Work in the last decade has begun to elucidate the roles of acetylcholine in epithelial cells and regulation of metazoan development (Angelini et al., 2004; Sánchez-Lazo et al., 2012; Strader et al., 2018; Wessler & Kirkpatrick, 2008b). The functions of AchRs in bilaterian cell-cell signaling range from regulating apoptosis and proliferation to regulating differentiation by modulating transcription (Maouche et al., 2009). Interestingly, the non-neuronal roles identified for AchRs are reminiscent of downstream effects of acetylcholine that have been identified in organisms lacking dedicated receptors suggesting that once receptors evolved they were incorporated into ancestral pathways to regulate cell biology in response to acetylcholine.

Two receptors have expression consistent with a role for acetylcholine in metamorphosis. *NvnAchRαJ* and *NvnAchRαK* are expressed in the apical organ of free swimming late stage planula (Figure 2O, S)(Sinigaglia et al., 2015). The apical organ is thought to regulate settlement and metamorphosis in some cnidarians. Acetylcholine has been found to be involved in the settlement and metamorphosis of bivalve free swimming larvae including *Pinctada maxima*, *Crassostrea gigas*, *Mytilus edulis*, and *Mytilus galloprovincialis* as well as within coral planula larvae (Sánchez-Lazo et al., 2012; Strader et al., 2018). In *Nematostella* the apical organ expression disappears once planula metamorphose to juvenile polyps indicating that acetylcholine may regulate the ability of the apical organ to promote metamorphosis within *Nematostella* (Figure 2P, T).

### Muscarinic receptors are unique to bilaterians

Acetylcholine has the ability to bind to both nicotinic acetylcholine receptors as well as muscarinic acetylcholine receptors. To date the domain necessary for GPCR signaling necessary for muscarinic acetylcholine receptors have not been identified within the genome of any cnidarian (Anctil, 2009; Kass-Simon & Pierobon, 2007). Pharmacological treatments performed within hydrozoans using the muscarinic antagonist, atropine, inhibited contractile bursting in *Hydra* and a hydrozoan jellyfish (Kass-Simon & Passano, 1978; Scemes & de-Freitas, 1989). However, the inability to identify muscarinic receptors within cnidarian genomes argues that the atropine responses are likely due to off target non-specific activation of non-muscarinic receptors. Our work confirms that *Nematostella* has no acetylcholine receptors that cluster with the bilaterian muscarinic acetylcholine receptors. The current data strongly argue that the muscarinic acetylcholine receptors are a bilaterian innovation.

### The evolution of acetylcholine signaling

The shared roles of acetylcholine in cell-cell communication found in bilaterians, metazoans that lack dedicated receptors, single celled organisms, and plants suggest that these roles represent the ancestral function of acteylcholine (Bamel, Gupta, & Gupta, 2016; Di Sansebastiano, Fornaciari, Barozzi, Piro, & Arru, 2014; Kawashima et al., 2007; Wessler & Kirkpatrick, 2008b). One hypothesis to explain the emergence of dedicated receptors is that as tissue and cell type complexity evolved, tissue specific regulation of these ancestral acetylcholine roles became crucial for precise regional regulation of gene expression, differentiation, and apoptosis. The evolution of acetylcholine receptors would have provided a mechanism to regulate when and which cells and/or tissue(s) would be competent to respond to acetylcholine as well as allow for tissue specific responses depending subunit composition of subunits within each receptor. An emerging area of developmental biology is trying to understand how changes in ion concentration and intracellular pH can influence specification and signaling. Ion channels provide a mechanism for cells to manipulate pH and intracellular ion concentration. Because the nicotinic receptors are ligand gated ion channels their evolution would have provided a mechanism to co-opt cholinergic signaling as a neuromodulator, which based on our data likely happened soon after emergence of the receptors.

To definitively determine the evolutionary tract of acetylcholine signaling the roles and mechanisms of acetylcholine activity in cell-cell signaling needs to be further studied in animals with and without dedicated receptors. Evidence for cholinergic signaling in early metazoans, plants, and bacteria without acetylcholine receptors has been identified; yet the mechanism is still unknown. Comparing how similar the molecular pathways by which acetylcholine regulates apoptosis, proliferation, cell differentiation, etc. in animals lacking receptors to how those pathways are regulated utilizing receptors is critical. This would confirm that signaling through the receptor represents maintenance of an ancestral function or a novel role for acetylcholine. For example, within bilaterians, non-neuronal acetylcholine activates a large number of downstream pathways, including protein kinase C (PKC) and mitogen-activated protein kinase (MAPK), to regulate non-neuronal functions (Cardinale, Nastrucci, Cesario, & Russo, 2012). Identifying similar pathway activation by acetylcholine in metazoans without acetylcholine receptors would indicate conservation in cholinergic cell-cell signaling. Further characterization of cholinergic signaling in *Nematostella* and other cnidarians is also necessary to confirm or reject the hypothesis that functions through receptors are conserved. In addition to comparing activity between animals with and without receptors an effort should be made to determine if acetylcholine is able to regulate the biology of cnidarians and bilaterians in a receptor independent fashion and if those putative activities resemble the functions identified in basally branching metazoans. The combined characterization of early metazoan cholinergic with known bilaterian signaling would then more accurately inform us on the evolution of acetylcholine signaling within all metazoans.

### Conclusions

Our data coupled with recent observations indicate that acetylcholine receptors function in cell-cell signaling and regulation of neuronal activity in cnidarians. Although, we and others observed receptor expression in muscles, the ability for lidocaine to block this acetylcholine induced tentacular contractions indicates that acetylcholine promotes contraction through an upstream neuron rather than acting directly on the myoepithelial cell. Our data and previously published results suggest that the cell-cell signaling and neuronal functions of acetylcholine signaling predates cnidarian-bilaterian divergence, but that the ability of acetylcholine to induce muscle contractions evolved within the bilaterians (Ahmed & Ali, 2016; Dani & Bertrand, 2007; Goldman & Staple, 1989; Jospin et al., 2009). We also hypothesize that evolution of acetylcholine receptors may have been selected for due their ability to control which cells, when cells, and how cells respond to acetylcholine, providing the added benefit of enhancing the ability to promote tissue/cell type specific responses to acetylcholine.

## Supplemental Data

**Supplemental Table 1: Alignment of the nicotinic acetylcholine receptors**

Protein alignment of confirmed bilaterian muscarinic and nicotinic acetylcholine receptors, glycine, and GABAergic receptors with *Nematostella* nicotinic acetylcholine receptors

**Supplemental Video 1: Treatment with 10mM acetylcholine 4X magnification**

Treatment with acetylcholine at 5 secs induces tentacle contractions in *Nematostella*.

**Supplemental Video2: Treatment with 10mM acetylcholine 10X magnification**

Treatment with acetylcholine at 10 secs induces tentacle contractions in *Nematostella*.

**Supplemental Video 3: Pretreatment with 500μM Mecamylamine and then 10mM acetylcholine 4X**

Treatment with acetylcholine at 8 secs after pretreatment with mecamylamine blocks acetylcholine induced tentacle contractions in *Nematostella*.

**Supplemental Video 4: Treatment with 10mM Nicotine**

Animals treated 10mM nicotine at 8 secs in *Nematostella* induced tentacle contractions.

**Supplemental Video 5: Pretreatment with 10mM Lidocaine and then 10mM acetylcholine 4X**

Animals were pretreated with 10mM lidocaine and then treated with acetylcholine at 8 secs showing loss of acetylcholine induced tentacle contractions in *Nematostella*.

**Supplemental Video 6: Treatment with water**

Animals were treated with water at 8 secs in *Nematostella*.

**Supplemental Table 2: Primers used for amplification**

## Abbreviations

nAchR: nicotinic acetylcholine receptor
mAchR: muscarinic acetylcholine receptor
hpf: hours post fertilization

## Authors’ Contributions

DZFG and ML conceived the idea and wrote the manuscript. DZFG carried out the collection and analysis of the data. All authors read and approved the final manuscript.

